# Separation slang – Laboratory mice use low-frequency call repertoire during physical separation

**DOI:** 10.1101/2025.09.26.678748

**Authors:** Daniel Breslav, Michal Wojcik, Ursula Koch, Thorsten Becker

## Abstract

The discovery of a diverse repertoire of ultrasonic vocalizations (USVs) sparked interest in understanding their role in mouse social behavior. Social communication in mice is not just vocal, but multimodal and occurs mostly in close proximity. Aiming to unravel the impact direct physical interaction has on the vocal communication of same-sex mouse dyads, we separated mice through a divider preventing direct physical interaction, but allowing visual, olfactory and some tactile interaction through holes. Separated dyads emitted a distinct call repertoire consisting mainly of calls in or just above the human audible range (but not squeaks) as well as Noisy calls, and only to a lesser degree of USVs. Increasing the possibility for direct interaction through larger holes in the divider led to an adaption of the call repertoire. The separation-induced call repertoire was neither affected by sex, nor was it mouse strain specific, even though differences in spectro-temporal parameters and call class proportion occurred. Lastly, buspirone treatment showed no observable effect, suggesting anxiety to not be the main driver underlying the separation-induced call repertoire. We show that separated same-sex mouse dyads predominantly emit a call repertoire that until now has only been observed in isolation or during aversive stimulation.

## Introduction

Social situations pose challenges for the individuals involved, as messages need to be composed appropriately and receivers need to be competent in deciphering said messages to avoid misunderstandings. Interestingly, in humans non-verbal signals outweigh verbal communication when conveying emotions (Mehrabian, 1971). Thus, multimodal communication is essential in social interactions. During courtship (Hanson and Hurley, 2012; Ronald et al., 2020), territorial same-sex encounters (Gourbal et al., 2004), or in the discrimination of stranger from familiar mice (de la Zerda et al., 2022), mice use a mix of olfactory, acoustic, and tactile signals to communicate with one another. The resurgence of interest in mouse vocalizations during the last years has culminated in a detailed description of the neural circuit underlying mouse vocalizations (Chen et al., 2021; MacDonald et al., 2024; J. Park et al., 2024; Tschida et al., 2019; Veerakumar et al., 2023; Ziobro et al., 2024). Mice possess a large repertoire of ultrasonic vocalizations (USVs), which has been divided into different categories ranging from eight (Holy and Guo, 2005) up to 22 (Sangiamo et al., 2020). Certain call categories were found to be emitted more frequently in certain social contexts. USVs containing frequency jumps and those featuring harmonic elements occur particularly during courtship and are emitted by male mice (Chabout et al., 2015; Klaus et al., 2025), while females emit audible squeaks concomitantly with defensive behaviors (Finton et al., 2017; Hood et al., 2023; Lupanova and Egorova, 2015). Furthermore, a call with decreasing pitch (i.e. downward spectrogram slope) precede dominant social behaviors (Sangiamo et al., 2020). Altogether, adult mice vocalize usually in very close proximity to one another (Neunuebel et al., 2015; Oliveira-Stahl et al., 2023; Sangiamo et al., 2020), and mice that were socially deprived show increased USV emission and eagerness to interact with conspecifics (Burke et al., 2018; Chabout et al., 2012). In most studies, where USVs were emitted, mice were in direct contact allowing for tactile and close-range olfactory cues. In contrast, mice housed in isolation conditions produced a novel broadband vocalization with noisy features and emitted only few USVs (Grimsley et al., 2016), indicating that environmental conditions strongly shape the vocal repertoire. This also suggests that mice may adapt their vocalization pattern in more complex environments where non-vocal communication is restricted or severely limited.

This study investigates how the prevention of direct physical interaction (e.g. social whisking) shapes the vocal communication of same-sex mouse dyads, aiming to improve our understanding of the information transferred by mouse calls. We show a distinct, separation-induced call repertoire consisting mainly of calls in the human audible range (but not squeaks) as well as noisy calls, and only to a lesser degree of USVs. Increasing the possibility for direct interaction leads to an adaption of the call repertoire. The separation-induced call repertoire is not affected by sex and is not mouse strain specific even though differences in spectro-temporal parameters and call class proportion occur, respectively. Lastly, buspirone treatment being without any observable effect suggests anxiety to not be the main driver underlying the separation-induced call repertoire.

## Results

### Same-sex mouse dyads produce calls of lower frequency when separated by a divider featuring holes

Mice exhibit a remarkable diversity of vocalizations that play key roles in social interactions, with their characteristics and types varying depending on contexts such as mating, territorial disputes (Arriaga and Jarvis, 2013; Gourbal et al., 2004; Neunuebel et al., 2015), and affective states (Grimsley et al., 2011). While most of the produced vocalizations occur in close physical proximity, where tactile stimuli strongly influence the vocal output (Lahvis et al., 2011; Moles et al., 2007; Oliveira-Stahl et al., 2023), it remains unclear how vocal communication is affected by the absence of direct physical contact.

To explore this question, we conducted experiments using 11 female pairs and 11 male pairs of adult FVB stranger mice in a controlled setting, designed to record vocalizations during physical separation. Each same-sex pair was placed in a sound-isolated environment with a transparent divider having holes in the bottom section, allowing for visual, olfactory and vocal interaction while preventing physical contact (Fig. 1A). Under these conditions, we observed that typical ultrasonic vocalizations (USVs) were infrequent. After 15 minutes of physical separation, the divider was removed, and recording continued for another 5 minutes as the dyads were united and able to physically interact (Fig. 1B). This served as a control to determine whether the removal of the divider would cause the mice to revert to their typical repertoire of USVs. We observed one exception, a female dyad emitted solely USVs with a high call rate (18.3 calls/min) regardless of being separated or being able to interact directly with one another. Hence, this dyad was excluded from the analysis. Given the low rate of detectable USVs during separation, we reanalyzed the sound recordings without any high-pass filter. This adjustment uncovered previously undetected vocalizations at lower frequencies. These vocalizations were unambiguously distinct from established USVs and occurred at markedly higher call rates across experiments.

**Fig. 1.**
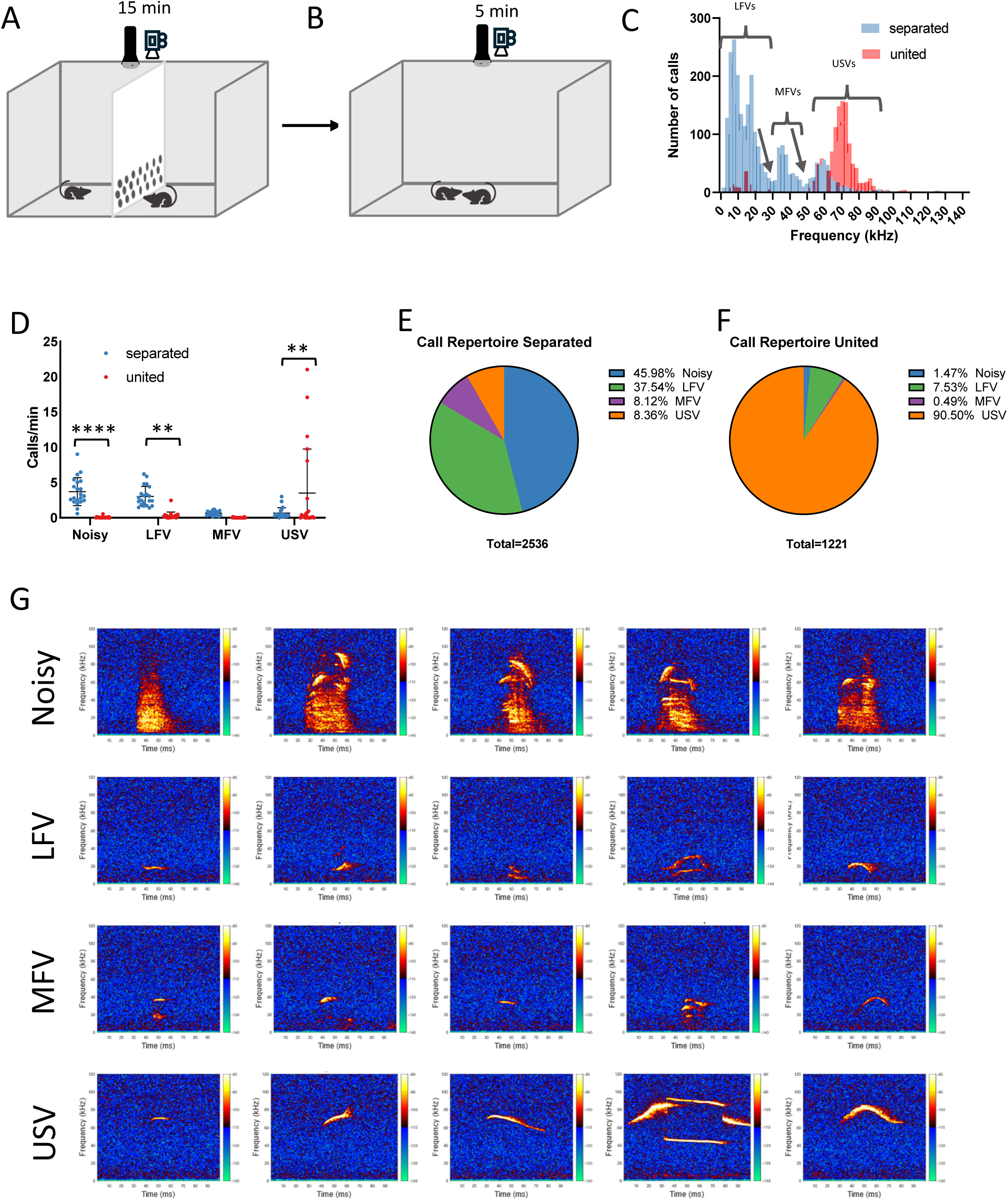

Analysis of peak frequency distribution of all vocalizations during both phases revealed a trimodal distribution for calls recorded during separation with two prominent troughs at 32 kHz and 50 kHz (Fig. 1C, blue histogram), resulting in the formation of three classes: Low-Frequency Vocalizations (LFVs; peak frequency <32 kHz), Middle-Frequency Vocalizations (MFVs; peak frequency 32–50 kHz), and USVs (peak frequency >50 kHz). Calls recorded during direct interaction showed peak frequencies nearly exclusively in the USV range, with only a few in the LFV range (Fig. 1C, red histogram). Additionally, a fourth class termed “Noisy” was identified. Noisy calls are characterized by a broad frequency spectrum (Fig. 1G, example spectrograms) and a distinctly warbled, noisy spectral appearance, and was distinguished based on their spectral morphology and large bandwidth (avg. 57.4 ± 6,74 kHz; Fig. 2D), rather than peak frequency. These vocalizations strongly resemble calls emitted during isolation as published by Grimsley and colleagues (2016).

**Fig. 2.**
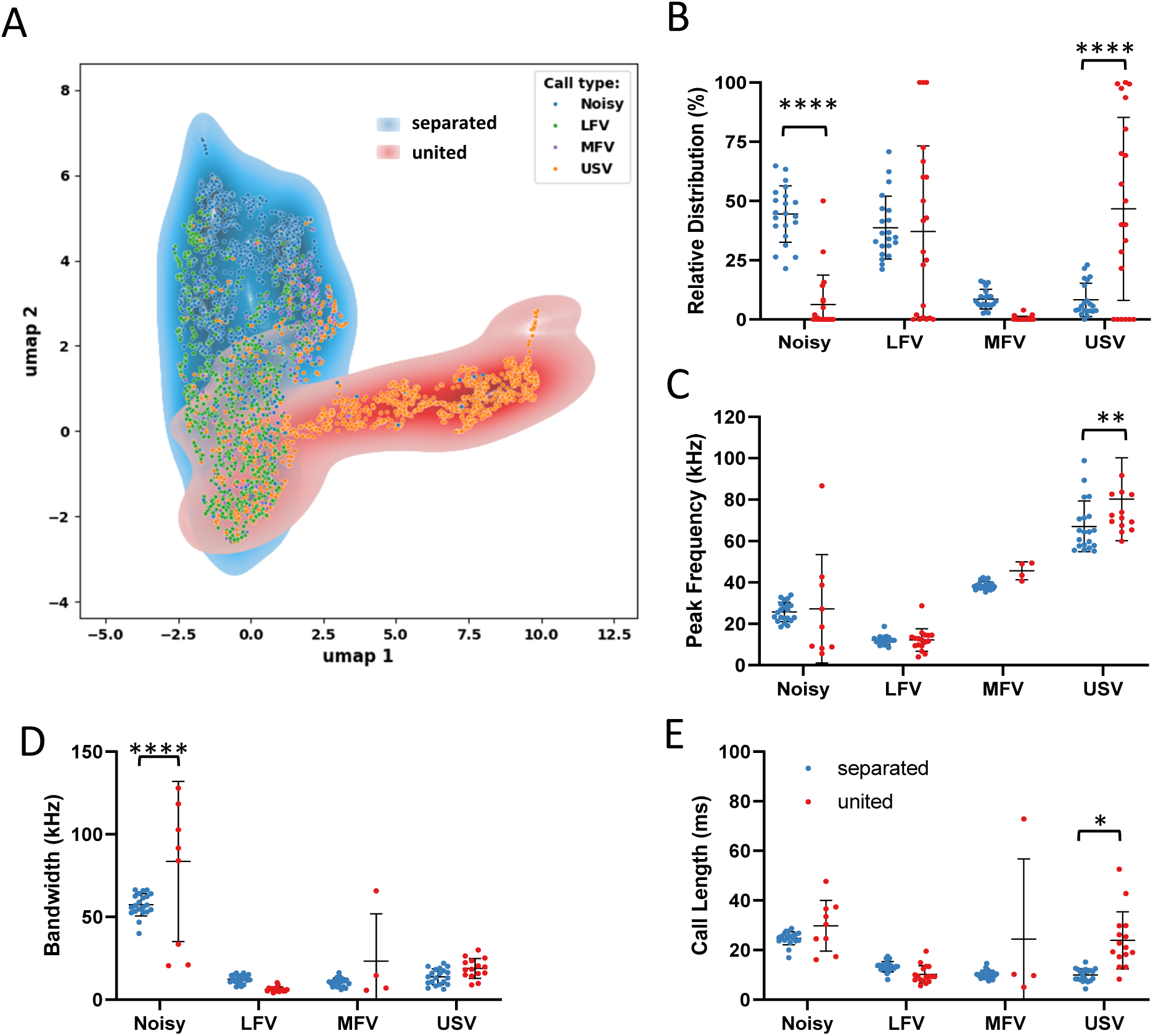

Separating two mice by means of a divider affected their call rate in a class-dependent manner (call class x separation: F_(3,80)_ = 14.80, P < 0.0001; Fig. 1D). While being separated, dyads emitted predominantly Noisy calls (3.7 ± 1.98 calls/min) and LFVs (3.0 ± 1.43 calls/min), however, when interacting freely call rates dropped to 0.2 ± 0.35 calls/min (P < 0.0001; Fig. 1D) and 0.9 ± 1.57 calls/min (P = 0.0019; Fig. 1D), respectively. The opposite was observed for USVs, call rate was low during separation and increased when they were united again (0.7 ± 0.32 calls/min vs. 10.5 ± 18.69 calls/min, P = 0.0012; Fig. 1D). MFVs were emitted at low call rates regardless of the context (0.7 ± 0.32 calls/min vs 0.1 ± 0.13 calls/min, P = 0.8689; Fig. 1D).

In total we recorded 2,536 vocalizations during separation, dyads emitted predominantly Noisy calls (45.98%) and LFVs (37.54%). MFVs (8.12%) and USVs (8.36%) accounted for the remaining calls (Fig. 1E). In contrast, of the 1,221 calls recorded during unification, over 90% were USVs and only 10% belonged to either Noisy calls (1.47%), LFVs (7.53%), or MFVs (0.49%) (Fig. 1F). Thus, same-sex dyads used a distinct call repertoire during physical separation compared to unification. Despite this striking difference between these call repertoires, all four call classes were present in both phases. Interestingly, like the widely known mouse USVs (Fig. 1G, USV), both LFVs and MFVs displayed diverse spectral morphologies (Fig. 1G, LFV, MFV). Surprisingly, even the spectrograms of Noisy calls displayed a large variety, with often different tonal components in both mid and high frequencies being visible (Fig. 1G, Noisy).

### Impact of separation on spectro-temporal call features

These findings prompted further investigation into which temporal and spectral properties of these calls, besides their peak frequency, could be used for further characterization. For instance, categories for mouse USVs have been formed based on their spectral morphology (Holy and Guo, 2005; Scattoni et al., 2009, 2008), raising the possibility for more refined categories of both LFV and MFV. Thus, we employed the convolutional neural network (CNN) VGG16 to perform call feature extraction based on the call spectrogram images resulting in 4096 different features. Dimensionality reduction to two dimensions by UMAP revealed two distinct clusters: one corresponding to mainly USVs (Fig. 2A, yellow dots) and a second cluster encompassing LFVs, MFVs, USVs and Noisy calls. Within this latter cluster, a finer substructure was observed, with Noisy calls (Fig. 2A, blue dots) segregating toward the north-eastern region, LFVs (Fig. 2A, green dots) in the south-western region, and MFVs (Fig. 2A, purple dots) in the eastern region while USVs were present throughout the cluster. Furthermore, the USV cluster consisted predominantly of USVs that were emitted during unification (Fig.2A yellow dots on dark red shade), which segregated from the other call types, even USVs emitted during separation (Fig. 2A yellow dots on blue shade).

In conclusion, firstly, unsupervised, spectrum-based mouse call segmentation using a VGG16 CNN revealed a large part of USVs emitted during unification were distinctly different from USVs emitted when separated. Secondly, LFVs, MFVs, and Noisy calls were not further divided into distinct subclusters. This has previously been shown for USVs (Goffinet et al., 2021; Sainburg et al., 2020). Therefore, we decided to refrain from further categorizing these classes for now.

While all same-sex dyads emitted calls during separation, the number of calls emitted varied considerably between individual dyads (range: 42 to 269 calls, avg. 121 calls). Also, when united, 19 out of 21 dyads emitted calls, ranging from a total of 2 calls up to 317 calls (avg. 64 calls). This high variance prompted us to investigate the relative distribution of each call class per dyad and separation condition. Similar to the call rates, the relative distribution of call classes, particularly, of Noisy and USV calls was affected by the separation (call class x separation: F_(3,80)_ = 25.29, P < 0.0001; Fig. 2B). While Noisy calls made up 44.5 ± 11.92% of calls during separation, once united they only accounted for 6.3 ± 12.44% of the dyads’ call repertoire (P < 0.0001; Fig. 2B). The opposite was observed for USVs, during separation dyads emitted USVs rarely (8.3 ± 7.06%), however, once the animals were able to interact directly USVs made up a substantial number of calls (46.7 ± 38.60%, P < 0.0001; Fig. 2B). While the absolute call rate of LFVs was significantly higher during separation compared to unification (Fig. 1D), the relative call type distribution did not significantly differ between events but displayed a huge disparity between subjects (38.7 ± 13.28% vs 37.1 ± 36.09%, P = 0.9984; Fig. 2B). Regarding MFV distribution, same-sex dyads emitted more MFVs during separation (8.6 ± 4.19%) than during unification (0.4 ± 0.98%), however, this difference was not statistically significant (Fig. 2B). Beyond the effect separation had on the call repertoire used by same-sex dyads, we observed a lower peak frequency of 67.1 ± 12.25 kHz in USVs emitted while mice were separated compared to 80.2 ± 20.0 kHz during unification (P = 0.0037; Fig. 2C). Also, the average duration of USVs was substantially longer when two mice interacted freely with one another (23.9 ± 11.54 ms vs 10.0 ± 2.59 ms, P = 0.0244; Fig. 2E). Nevertheless, physical separation did not affect the peak frequencies or call lengths of the other call classes. Spectral properties peak frequency might not be suitable to describe Noisy calls. Instead, bandwidth seemed to be a much more fitting parameter. Indeed, we found the bandwidth of Noisy calls to be quite consistently between dyads separated by the divider (57.4 ± 6.74 kHz, Fig. 2D), while the bandwidth of Noisy calls emitted during free interaction varied more substantially (83.6 ± 48.43 kHz, P < 0.0001; Fig. 2D). It is noteworthy that not all dyads emitted Noisy calls when the two mice could interact freely.

Together, these results demonstrate that physical separation through a divider evoked a call repertoire containing predominantly Noisy calls and LFVs and only to a minor part MFVs and USVs, while freely interacting mice emitted mainly USVs. Furthermore, the few USVs emitted during separation were shorter and of lower peak frequency than those produced during unification. During physical separation Noisy calls exhibited a lower bandwidth than during direct physical interaction.

### Temporal organization of separation-induced calls

We observed a substantial number of calls from all four classes occurring in groups or bouts. These bouts could contain solely calls from one class, i.e. Noisy, LFV, USV (Fig. 3A-C), but could also contain a mix of different call classes (Fig. 3D). Analysis of the inter-call interval (ICI) distributions of all four call classes revealed a peak at an ICI of about 90 ms during both separation (Fig. 3E-H, blue histograms) and unification (Fig. 3E-H, red histograms). All calls with an ICI of 140.6 ms or lower (the trough following the 90 ms peak) were considered to be emitted in bouts. When separated, dyads emitted Noisy calls predominantly in bouts (77.2%), while every second LFV was also embedded into a bout (54.3 %). Even though fewer MFVs and USVs were produced than both Noisy and LFV calls during separation, 44.1% of MFVs and 43.1% of USVs were emitted in bouts. Once mice were allowed to interact freely, the majority of calls recorded were USVs (87.1% in bouts), while 43.1% of LFVs were emitted in bouts. Of the few MFVs and Noisy calls emitted 50% and 17.7% were part of a bout, respectively.

**Fig. 3.**
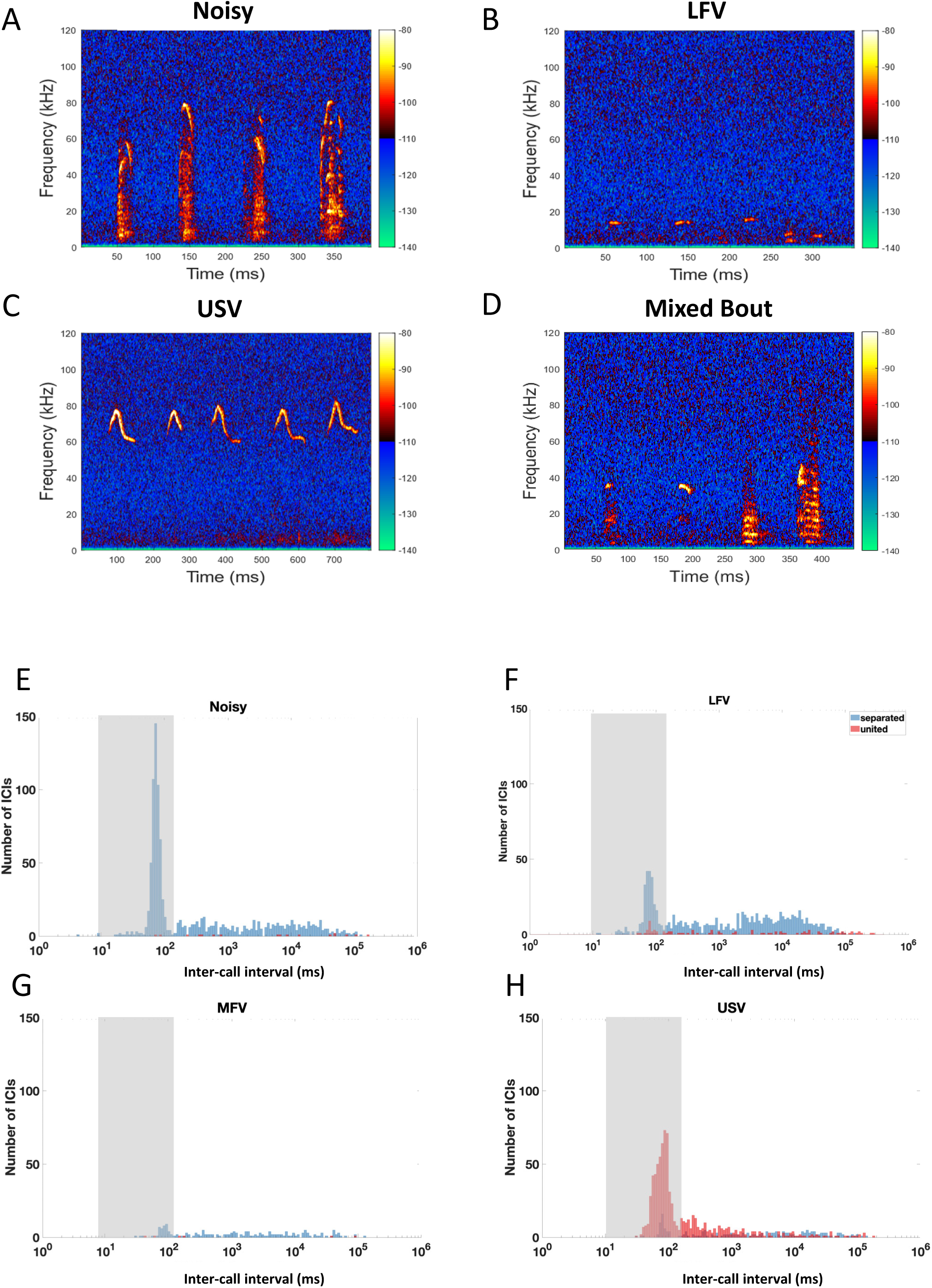

Same-sex mouse dyads’ call repertoire displayed a condition-dependent temporal organization. While separated, about 80% of Noisy calls were emitted in bouts. This was the case also for about 40 to 50% of LFVs, MFVs, and USVs. In contrast, in freely interacting mice about 90% of USVs emitted were structured in bouts. Our findings suggest that vocal communication in mice is highly condition-dependent, with different vocalization strategies emerging in response to physical separation.

### Male and female same-sex dyads emitted similar separation-induced call repertoire, but with different spectro-temporal features

Building on these findings, we next sought to investigate whether the observed differences were influenced by the sex of the mice. The call repertoire used by male same-sex dyads did not differ from that of female same-sex dyads (Fig. 4A -D). However, separated female dyads tended to emit calls at a higher rate (9.4 ± 3.79 calls/min) compared to separated male dyads (6.8 ± 2.33 calls/min, t_(19)_ = 1.907, P = 0.0718; Fig. 4E). Furthermore, female dyad calls exhibited a higher bandwidth (36.3 ± 4.80 kHz) and were longer (19.3 ± 2.01 ms) compared to male dyad calls (28.8 ± 6.66 kHz, t_(19)_ = 2.931, P = 0.0086; Fig. 4G) (16.7 ± 2.64 ms; t_(19)_ = 2.548, P = 0.0197; Fig. 4H). To verify whether these observations were context-dependent, we examined the vocalization repertoires in opposite sex pairs separated by the perforated divider. Mice are known to emit a large amount of USVs during courtship (Hanson and Hurley, 2012; Wang et al., 2008), even when separated by a divider male-female dyads emitted a substantial amount of USVs (Hood et al., 2023; Hood and Hurley, 2023). Similarly, in our separation condition mixed-sex dyads emitted predominantly USVs (80.0%; Fig. 4 - Suppl. 1). This call repertoire exhibited a close resemblance to the repertoire observed during unification (Fig. 1F), though not identical, as separated mixed-sex dyads produced a larger proportion of both Noisy calls (mixed-sex: 6.60 % vs same-sex: 1,47%) and MFVs (mixed-sex: 4.58% vs same-sex: 0.49%) compared to separated same-sex dyads. It contrasted starkly with the call repertoire of same-sex dyads that consisted mainly of Noisy calls and LFVs (Fig. 1E).

**Fig. 4.**
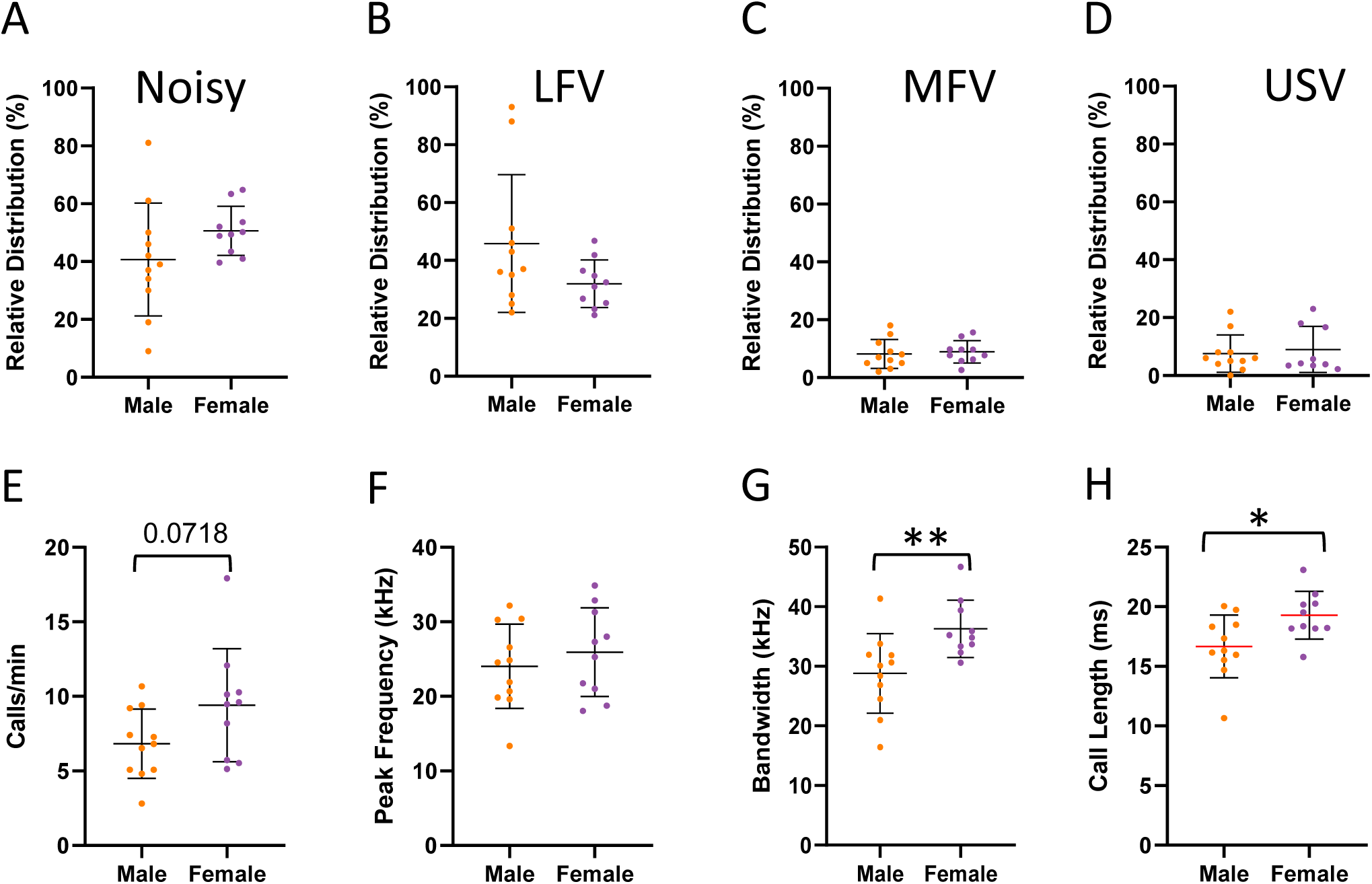

Summing up, the call repertoire and call features did not show any major differences between male and female same-sex dyads. Mixed-sex dyads emitted predominantly USVs even when separated.

### Separated same-sex dyads were most likely to vocalize in close proximity while facing one another

To investigate the relationship between vocal activity and the relative position of mice to each other, we tracked mouse body parts using DeepLabcut. When separated calls were detected at nearly all snout-to-snout distances throughout the experiment in both male and female dyads. However, despite the obstruction of direct physical contact by the divider, mice emitted vocalizations particularly often in very close proximity, i.e. a snout-to-snout distance closer than one mouse body length (length: 200 px ≙ 6 cm, Fig. 5 C1, C2, D1, D2 grey bar). Separated male dyads emitted predominantly Noisy calls in very close proximity, while separated female dyads displayed a more varied call repertoire in close proximity consisting mainly of Noisy calls and LFVs, but to a lower extent also of MFVs and USVs (Fig. 5 C1, D1 colored lines in grey bar). After allowing direct physical interaction by removing the divider, nearly all vocalizations recorded were produced within a distance of two body lengths (i.e. 0 to 400 px) in both male and female dyads. When united female dyads produced nearly exclusively USVs, whereas male dyads also emitted LFVs in close proximity (Fig. 5 C2, D2). Incorporating the snout-to-snout angle (Fig. 5 A) into the analysis revealed a positive correlation between snout-to-snout distance and angle (table 1). When separated dyads were facing one another when emitting calls in close proximity (at 0 to 100 px distance, 0 to 60° snout-to-snout angles, Fig. 5 E1, F1), while they were less likely to face one another the further they were apart. After the divider was removed, same-sex dyads displayed normal social behaviors, such as anogenital sniffing and following/pursuit behavior, which was reflected by the larger number of calls emitted at greater snout-to-snout angles (Fig. 5 E2, F2, 50° and upward). To investigate this further, we calculated the snout-to-tail base distance and angle between the two mice (Fig. 5 B). Anogential sniffng is represented by small snout-to-tail base distances (200 px or less), while the large range of snout-to-tail base angles represents the body position relative to one another. 0° signifies the two mice being in-line with one mouse’s snout ‘facing’ the other mouse’s tail base, while at a snout-to-tail base angle of 175° the two mice were facing in opposite direction (Fig. 5E2, F2 inset scatter plots). Calls emitted during following/pursuit behavior were reflected by larger snout-to-tail base distances (200 px and up) and smaller snout-to-tail base angles (0 – 50°, Fig. 5 E2, F2 inset scatter plots). It has to be noted that both snout-to-tail base distance and angle alone are not suffcient regarding the discrimination of whether a behavior constitutes in-line anogenital sniffing or the beginning of following/pursuit behavior, as this was not the goal of this analysis.

**Fig. 5.**
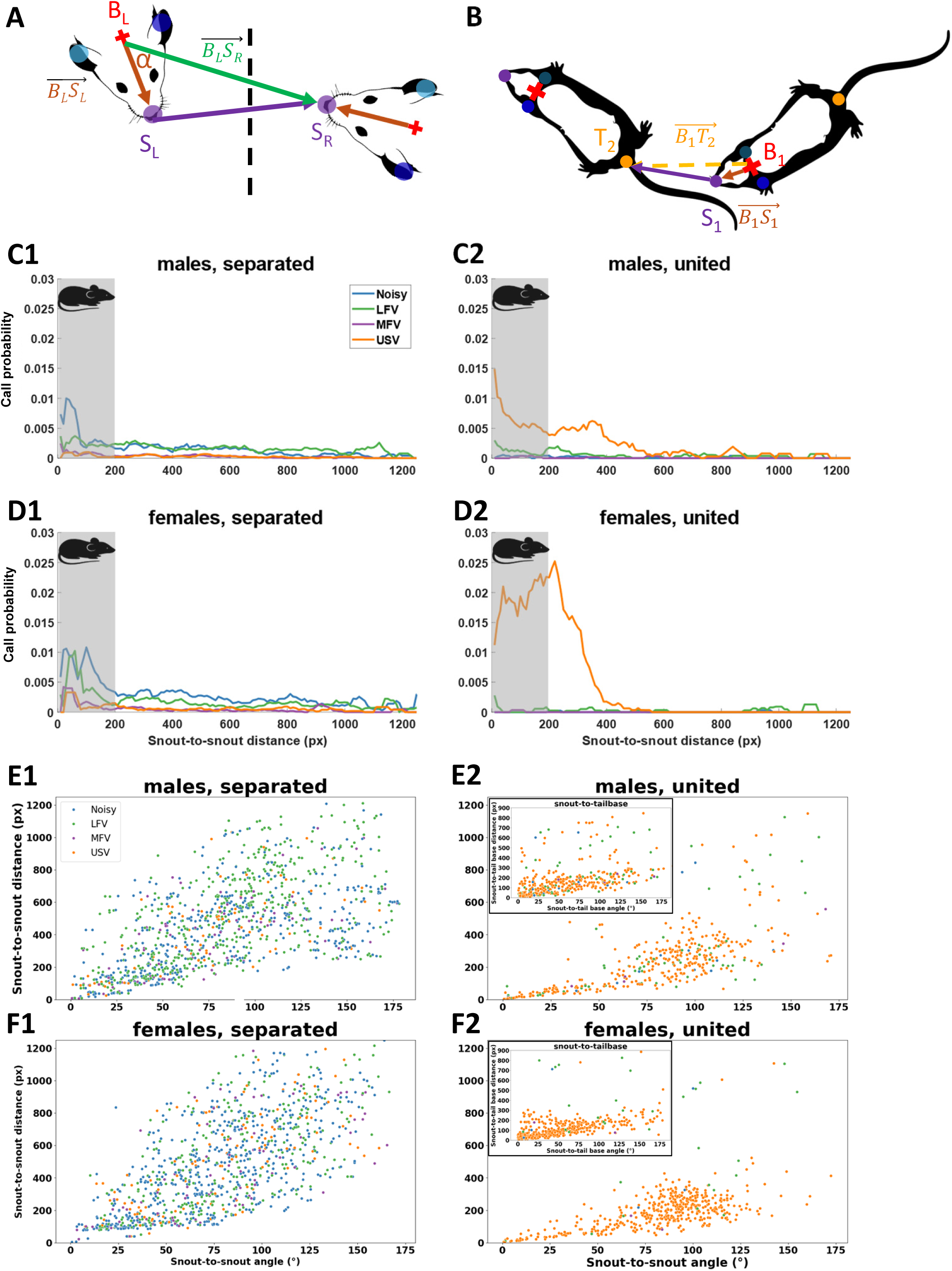

**Table 1.**
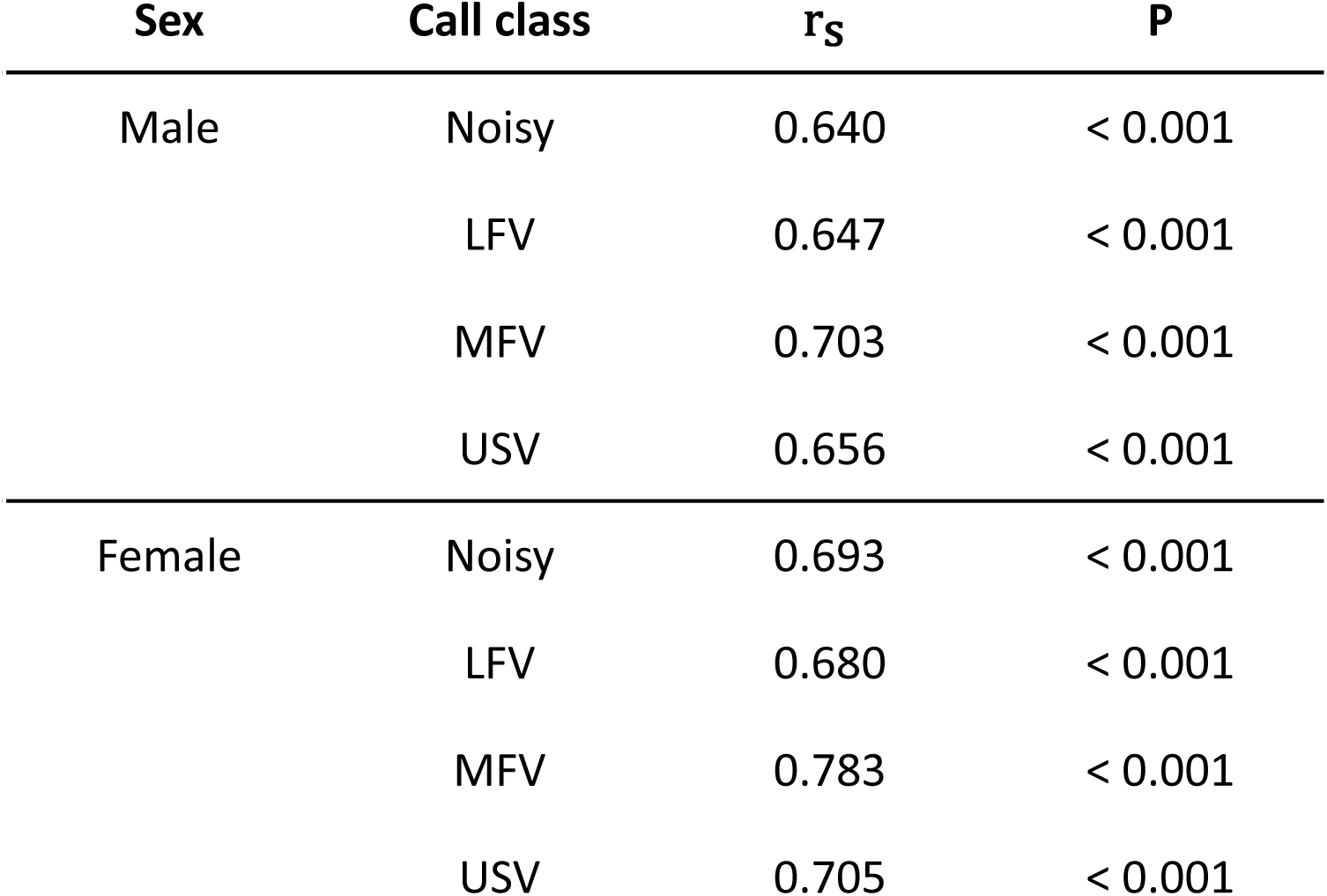
Spearman correlation between snout-to-snout distance (px) and snout-to-snout angle (°) for separated same-sex mouse dyads.

In summary, while direct physical interaction was obstructed by the divider, the dyad was most likely to emit calls in close proximity while facing one another at the divider. During unobstructed direct physical interaction both male and female same-sex dyads emitted mostly USVs. The emission of these USVs occurred mainly during anogenital sniffing and following/pursuit behavior. These observations suggest that two same-sex mice resort to a form of directed, vocal communication, while being prevented from direct physical interaction.

### Distinct separation-induced call repertoires and spectro-temporal call properties in different mouse strains

To investigate whether these newly identified calls are caused by a specific genetic background, we analyzed separation-induced calls in two additional mouse strains with different genetic backgrounds (Fig.6A). CBA mice were selected for their suitability in auditory research, as they maintain relatively stable hearing thresholds over age (Wu et al., 2019, Ohlemiller et al. 2010). In contrast, C57BL/6J (B6J) mice are widely used in biomedical research including vocalization behavior studies, despite their early onset, progressive hearing loss (Henry & Lepkowski, 1978). All tested strains produced the full repertoire of call classes (Fig. 6 B – D), however, strain-specific differences emerged.

**Fig. 6.**
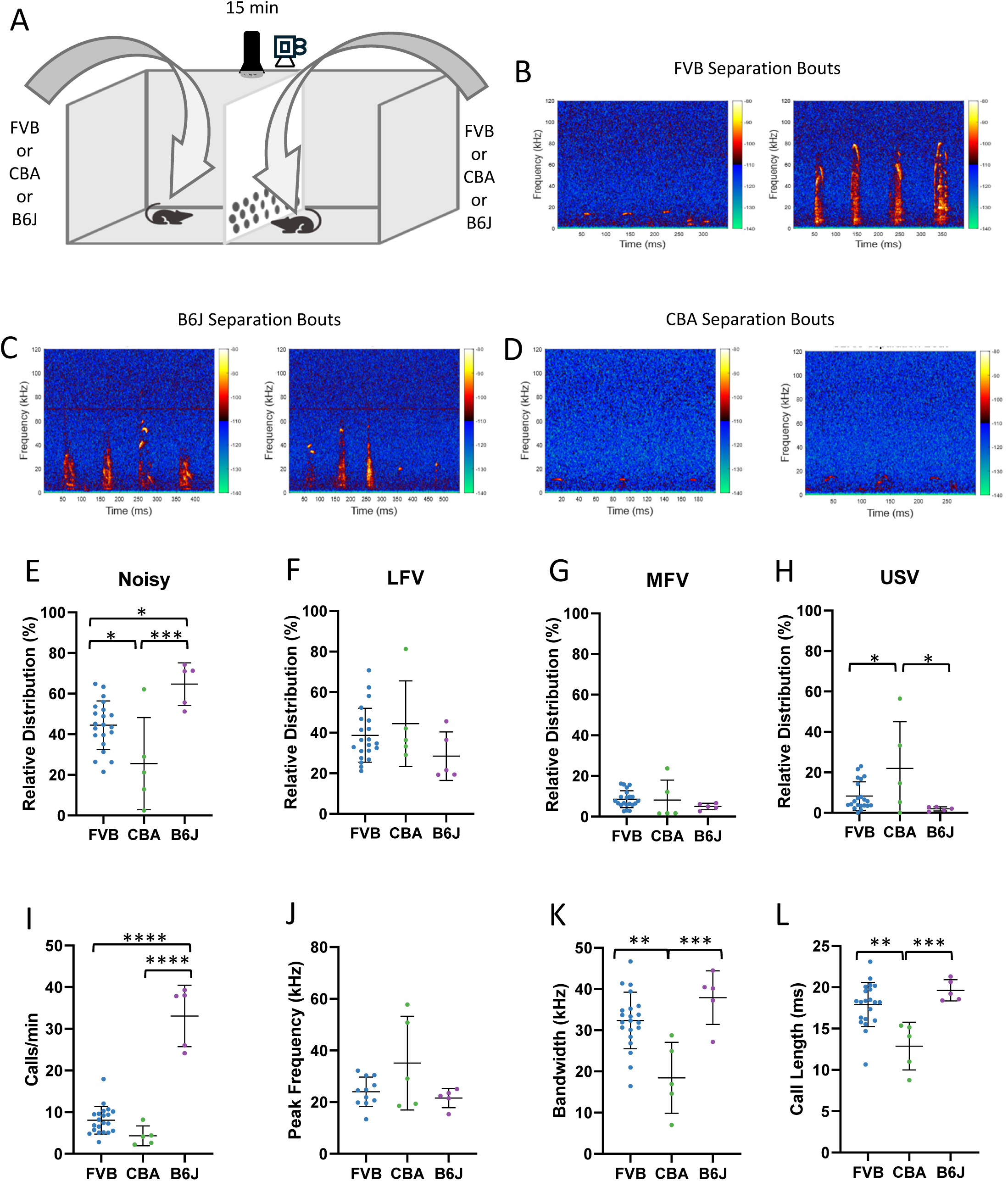

With a ratio of 25.5 ± 22.7%, separated same-sex CBA dyads emitted fewer Noisy calls than both FVB (44.5 ± 11.9%) and B6J mice (64.7 ± 10.5%) (FVB vs. CBA P = 0.0268, FVB vs B6J P = 0.0172, CBA vs. B6J P = 0.0003; Fig. 6 E). Conversely, CBA mice produced the highest proportion of USVs (21.9 ± 23.1%), significantly exceeding both B6J (1.9 ± 1.0%) and FVB (8.3 ± 7.1%) mice (FVB vs. CBA P = 0.0384; CBA vs. B6J P = 0.0153; Fig. 6 H).

B6J mice vocalized at a significantly higher rate (33 ± 7.4 calls/min), producing more calls per minute than both CBA (4 ± 2.4 calls/min) and FVB (8 ± 3.3 calls/min) mice (FVB vs. B6J P < 0.0001; CBA vs. B6J P < 0.0001; Fig. 6 I). Despite their higher call rates, B6J mice did not differ significantly from FVB mice in bandwidth or call duration (Fig. 6 K, L). In contrast, CBA mice displayed a markedly narrower bandwidth (18.5 ± 8.6 kHz) compared to FVB (32.4 ± 6.9 kHz) and B6J mice (37.9 ± 6.5 kHz; Fig. 4K, FVB vs. CBA P = 0.0014; CBA vs. B6J P = 0.0005; Fig. 6 K). CBA mice also displayed significantly shorter call durations (12.9 ± 2.9 ms) compared to both FVB (17.9 ± 2.7 ms) and B6J (19.6 ± 1.3 ms) mice (FVB vs. CBA P = 0.0013, CBA vs. B6J P = 0.0007, Fig. 6 L).

These findings demonstrate that genetic background profoundly shapes the vocal communication repertoire in mice, influencing not only the proportional distribution of call classes but also the production rates and spectral properties of calls.

### Varying sensory interaction during separation affects call repertoire and call features

We examined how different divider hole sizes (small, large or none) influence the call repertoire due to varying degrees of visual, olfactory, and putatively tactile interaction. Larger diameter (1 cm) of the holes resulted in a significant change of the vocal repertoire compared to that observed with the small holes (0.5 cm) or a solid divider (Fig. 7 B, C, D). Large holes led to a markedly lower proportion of Noisy calls (22.2 ± 10.6%), compared to small (44.5 ± 11.9%) and none (44.5 ±16.0%) (F_(2,34)_ = 9.758, P = 0.0004; Fig. 7 E). In turn, large holes were associated with a significantly higher proportion of LFVs (75.6 ± 11.1%) compared to small (38.7 ± 13.3%) and none (42.1 ± 12.4%) (F_(2,34)_ = 25.39, P < 0.0001; Fig. 7 F). Both MFVs and USVs accounted for about 1 % of the calls when mice were separated by a divider with large holes, while both MFVs and USVs made up about 9% and 7% when the hole diameter was small or the divider was solid, respectively (MFVs: F_(2,34)_ = 12.53, P < 0.0001; Fig. 7 G, USVs: H_(2)_ = 14.92, P = 0.0006; Fig. 7 H).

**Fig. 7.**
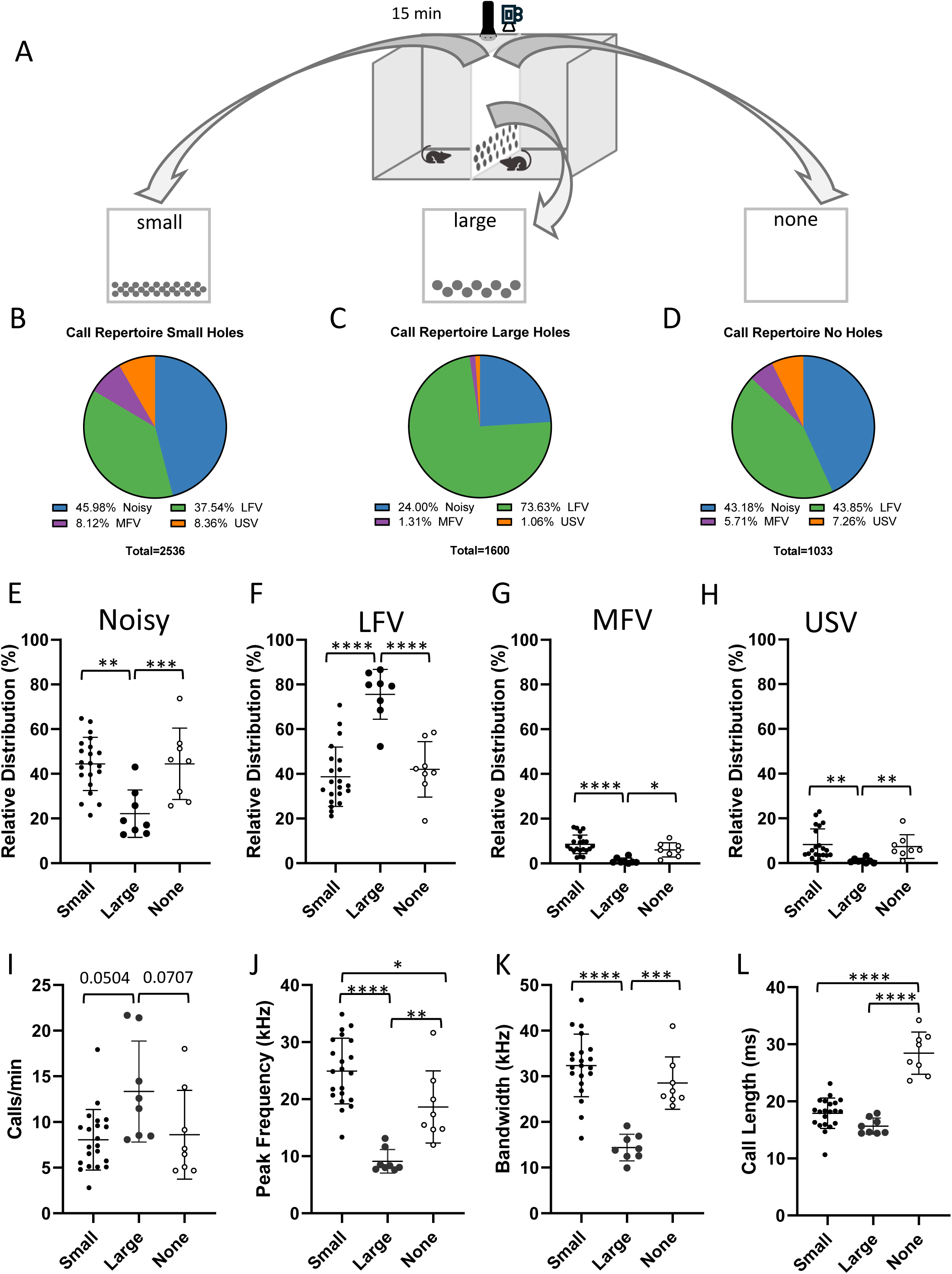

Also, the spectral and temporal call properties varied significantly with hole diameter. The hole diameter affected the call rate in separated dyads (χ^2^(3) = 6.785, P = 0.0336; Fig. 7 I). When the divider featured large holes, mice emitted about 13 calls/min compared to both 8 calls/min with small holes and 9 calls/min with a solid divider (large vs. small P = 0.0504; large vs none P = 0.0707, Fig. 7 I). Peak frequency was also impacted by the hole diameter (F_(2,34)_ = 25.86, P < 0.0001; Fig. 7 J). The lowest average peak frequency of about 9 kHz was observed with large holes, followed by an average peak frequency of about 19 kHz when the divider was not featuring any holes. Lastly, the highest average peak frequency of about 25 kHz was observed with small holes (small vs. large P < 0.0001, small vs. none P=0.0199, large vs. none P = 0.0031; Fig. 7 J). Similarly, hole diameter impacted the average call bandwidth (F_(2,34)_ = 25.92, P < 0.0001; Fig. 7 K), with the narrowest (∼ 15 kHz) occurring with large holes and similar average bandwidths with small (∼ 32 kHz) or no holes (∼ 29 kHz; small vs. large, P < 0.0001, large vs. none, P = 0.0001). Furthermore, the presence of holes in the divider impacted the average call length (F_(2,34)_ = 54.62, P < 0.0001; Fig. 7 L), with shorter call length occurring when the divider featured either small (∼ 18 ms) or large holes (∼ 16 ms) compared to an average call length of about 28 ms when the divider was solid (small vs. none P < 0.0001, large vs. none < 0.0001).

These findings highlight the impact of varying sensory access on both call repertoire, production rate and acoustic features of mouse vocalizations. Larger holes that allowed more direct somatosensory, visual, and olfactory interaction between the two mice resulted in more calls with lower peak frequency and smaller bandwidth, while the solid divider increased call duration. This is somewhat surprising, as the separation-induced emission of LFVs would have been expected to be mitigated by increased sensory interaction.

### Buspirone Treatment Reduces Anxiety but Does Not Affect Low-Frequency Vocalizations in Mice, Indicating Anxiety Is Not the Primary Cause

Low-frequency vocalizations, as observed in our experiments, have previously been reported primarily in aversive contexts involving restraint or low temperature environments (Grimsley et al., 2016; Yamauchi et al., 2022). Based on this, we aimed to investigate whether treatment with the anxiolytic buspirone would alter the call repertoire of separated mice to vocalizations of higher peak frequency and diminish the use of low-frequency vocalizations. Buspirone was shown to cause an anxiolytic effect in various behavior tests without the sedative effect known from benzodiazepines such as diazepam (Onofre-Campos et al. 2023). To test this, we compared vocalizations emitted after buspirone treatment to a control vehicle (saline) condition. Each mouse received both vehicle and buspirone treatments in a counterbalanced order, with a recovery period between sessions to control for potential carryover effects.

Vocal behavior of female same-sex dyads was impacted by the repeated exposure to the apparatus, but not the pharmacological treatment (Fig. 8 – Suppl. 1), thus, female dyads were excluded from the analysis. In contrast, vocal behavior of male dyads appeared unaffected by the repeated exposure. Interestingly, treatment with buspirone did not seem to affect the separation-induced call repertoire, compared to vehicle treatment (Fig. 8 B, C). However, including data of previously used untreated male same-sex dyads (Fig. 4) revealed untreated mice appeared to have emitted fewer Noisy calls, more MFV and USV calls (Fig. 8 D). The quantitative analysis of anxiolytic treatment effects on the four call classes supported this observation. Buspirone administration had no effect on the call distribution compared to vehicle treatment (Fig. 8 E - H). Compared to untreated mice, however, vehicle treated animals emitted more Noisy calls (52.9 ± 13.73 % vs 38.8 ± 12.11 %, t_(14)_ = 2.064, P = 0.0580), fewer MFVs (3.2 ± 0.89% vs 8.2 ± 4.62%, t_(14)_ = 2.368, P = 0.0328; Fig. 8 G) and USVs (2.8 ± 1.02 % vs 8.1 ± 6.83 %, U = 9, n_1_ = 5, n_2_ = 11, P =0.0380; Fig. 8 H).

**Fig. 8.**
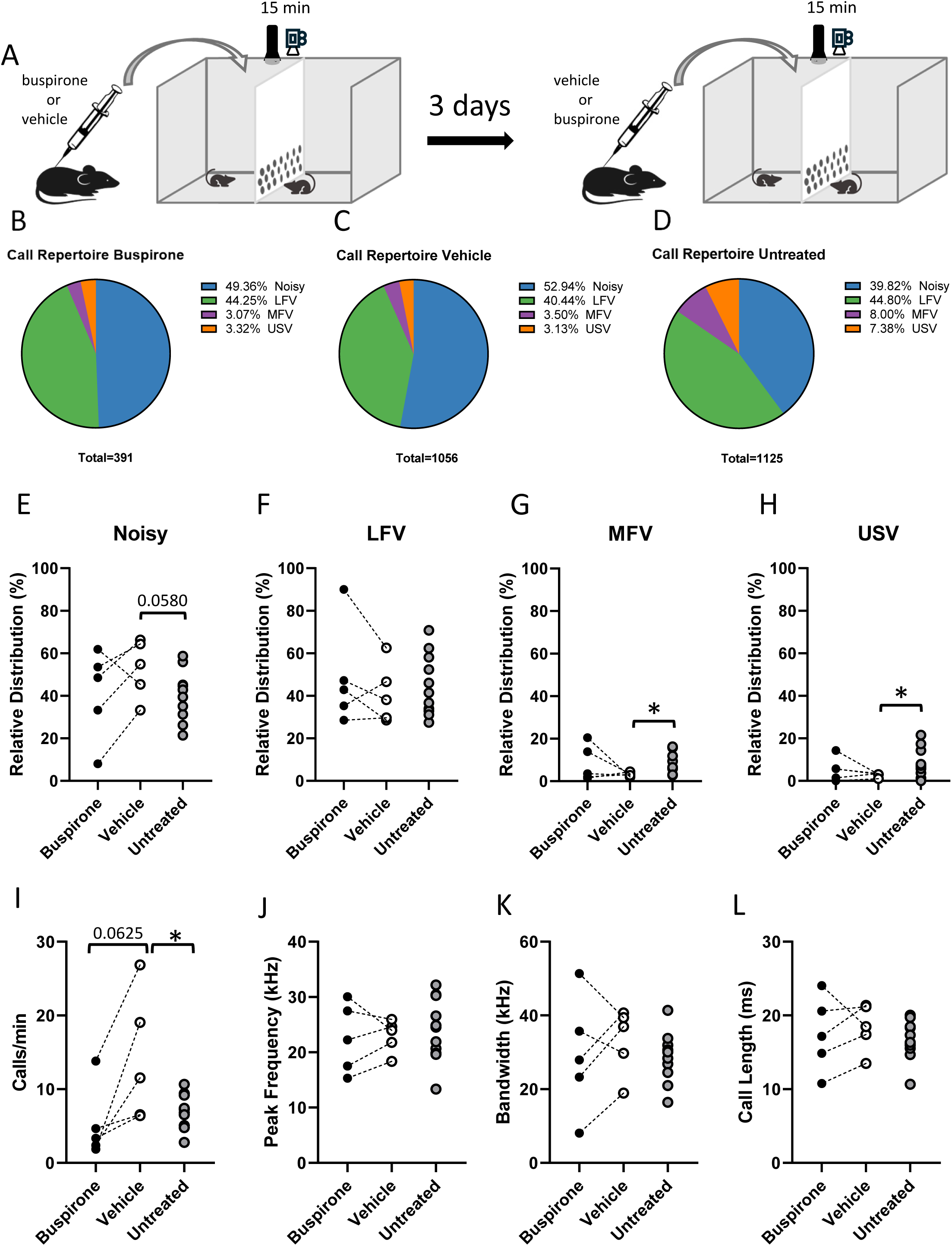

When investigating the spectro-temporal parameters, we did not observe any effect of either buspirone or vehicle treatment on peak frequency, bandwidth or call length. However, following the vehicle administration dyads emitted about 14 calls/min, while the same dyads emitted about 5 calls/min after buspirone treatment (W = 15.00, P = 0.0625; Fig. 8 I). The call rate of vehicle treated male dyads also exceeded that of untreated male dyads (∼7 calls/min; t_(14)_ = 2.637, P = 0.0195; Fig. 8 I).

Together, these results suggest that the low-frequency call classes observed during separation are unlikely to be driven primarily by anxiety, as the administration of the anxiolytic buspirone did not affect the call repertoire. However, the increased call rate observed in vehicle-treated mice indicates that the injection process itself may induce mild discomfort, which is mitigated by buspirone treatment.

## Discussion

The prevention of direct physical interaction of stranger same-sex mouse dyads through a divider revealed a separation-induced call repertoire. This repertoire consisted of calls in three frequency bands 1 – 32 kHz (low frequency vocalizations, LFV), 32 – 50 kHz (middle frequency vocalizations, MFV), and > 50 kHz (ultrasonic vocalizations, USV) and an additional class of Noisy calls, which is characterized by their broad bandwidth and warbled spectral appearance. Noisy calls and LFVs dominated the separation-induced call repertoire, while MFVs and USVs only accounted for a small portion. Both male and female FVB dyads used the same call repertoire, however, the females’ calls bandwidth being larger and the duration longer than that of the males. Furthermore, we observed a tendency towards a higher call rate in female dyads. Three different mouse lines commonly used in vocalization experiments were able to produce all four call classes, but emitted them at different proportions during separation. When separated the two mice were most likely to emit calls when they were less than one body length apart, facing one another. Once united the same mice would display normal social behavior including anogenital sniffing, pursuit and fleeing behavior. They adjusted their vocal repertoire according to the degree of direct interaction. Both dividers with no holes and with small holes resulted in a very similar call repertoire, while dyads encountering large-hole dividers emitted a vocal repertoire predominantly consisting of LFVs with an elevated call rate and fewer Noisy calls. Overall, the separation-induced call repertoire seemed not to be anxiety-related, as the anxiolytic buspirone had no effect on the repertoire, call rate, and spectro-temporal properties.

### Noisy mouse calls

#### Noise in mouse calls – feature, not artifact

Noisy calls were one of the two dominant call classes in the separation-induced call repertoire in our study. They pose a particular challenge with respect to their spectrogram-based identification as calls and differentiation from noise of non-vocal origin. However, we found evidence suggesting mice emit Noisy calls voluntarily in a context-dependent manner:

Firstly, 63.7% of Noisy call spectrograms contained a clearly visible tonal component. This could be explained by the use of the two distinct vocalization production mechanisms, vocal cord vibration and whistle, which are found in USV-producing rodent species (Mahrt et al., 2016; J. Park et al., 2024; Pasch et al., 2017; Riede et al., 2022; Roberts, 1975).

Secondly, most Noisy calls (77.2%) were emitted in bouts. In fact, mice have been shown to emit the majority of their USVs in bouts, both in courtship (Castellucci et al., 2018; Chabout et al., 2015) and in same-sex context (Hoier et al., 2016). The emission of vocalizations in bouts has also been described in other rodent species such as *Scotinomys teguina*, *Onychomys sp.*, and *Peromyscus sp.* (Okobi et al., 2019; Pasch et al., 2017; Riede et al., 2022). In our study, bouts consisting purely of Noisy calls were temporally highly regular with a median ICI of around 80 ms, suggesting that one call was emitted during one exhalation (Sirotin et al., 2014). This aligns with the ICI durations described for USVs produced in bouts (Castellucci et al., 2018; Chabout et al., 2015).

Thirdly, changing context parameters such as the diameter of holes in the divider resulted in a call repertoire containing considerably fewer Noisy calls meaning mice adapted their separation call repertoire as a consequence of the increased degree of sensory exchange through the divider.

#### Attracting attention through Noisy calls

Based on the observations that Noisy calls make up 43% of all calls in isolated male mice in an open field and Noisy calls spanning nearly the entire mouse hearing range, a role as seeking calls has been suggested (Grimsley et al., 2016). Adult CBA/CaJ mice have been reported to emit Noisy calls which spectrograms have a warbled, noisy, broadband appearance likely corresponding to “deterministic chaotic elements” (Grimsley et al., 2011), much like the spectrograms of our Noisy calls. Indeed, non-linear phenomena (NLP), such as biphonations or deterministic chaos, are a common component in mammalian vocalizations and are in some species associated with a heightened arousal state, e.g. in infant elephants (Stoeger et al., 2011), big brown bats (Gadziola et al., 2012), and vervet monkeys (Mercier et al., 2019). Furthermore, NLP in vocalizations have been shown to “grab” the listener’s attention in red deer and koalas (Charlton et al., 2017; Massenet et al., 2025; Reby and Charlton, 2012). Thus, the Noisy calls emitted in our experiments might serve to attract the attention of conspecifics in mice, too.

In our study, facilitated exchange of visual cues and somatosensory interaction through large holes in the divider (e.g., social whisking) reduced the proportion of Noisy calls. Whisking, together with both auditory and olfactory interactions between same-sex conspecifics are necessary in discriminating a familiar from an unfamiliar conspecific (de la Zerda et al., 2022). Indeed, in a resident-intruder context Noisy calls are being emitted during the social whisking preceding fight onset (Gourbal et al., 2004). A temporal coordination between whisking and USV emission has been shown in rats with the concomitant emission of a particular call repertoire at an increased call rate while in whisker-to-whisker contact. Not only the USV emission is linked to whisking in rats, but during whisker-to-whisker contact the response of regularly spiking neurons in primary auditory cortex to USVs is modulated (Rao et al., 2014).

Our data show that Noisy calls were particularly prevalent when direct physical interaction was prevented or strongly restricted and both mice were in close proximity at the divider. Facilitating direct physical interaction through large holes in the divider resulted in a reduction of Noisy call emission, while unrestricted physical interaction saw a cessation of Noisy call emission. These findings, together with the broad bandwidth and warbled spectral properties, support the notion that Noisy calls serve to attract conspecifics’ attention in situations where other forms of direct physical interaction are hampered or impeded.

### Putative communicational role of LFVs beyond the vocal expression of negative affective states

Besides Noisy calls, the separation-induced call repertoire was dominated by LFVs, which presented with an average peak frequency of about 12 kHz and spectrograms mostly without harmonics. These LFVs are being produced by the same neural circuit that gives rise to USVs, encompassing neurons in the dorsolateral periaquaeductal gray (PAG) projecting to neurotensin-positive neurons in nucleus retroambiguus (*Nts*^+^ RAm neurons) (Veerakumar et al., 2023), from which brain stem motor neurons controlling the vocal cords are being innervated (Chen et al., 2021; J. Park et al., 2024; Tschida et al., 2019; Veerakumar et al., 2023; Ziobro et al., 2024). While neuronal activity of USV-eliciting PAG neurons scales the amount of USVs and their loudness (Chen et al., 2021), *Nts*^+^ RAm neuron activity scales the dominant frequency of mouse calls (Veerakumar et al., 2023). This dominant frequency scaling constitutes a stepwise transformation induced by crossing a spike frequency threshold. Optogenetic stimulation of *Nts*^+^ RAm neurons in the range of 15 to 20 Hz reliably evoked the emission of LFVs, while stimulation frequencies > 25 Hz led to the emission of USVs (Veerakumar et al., 2023).

LFVs (corresponding to MFVs published by Grimsley et al. 2016) have been reported to vocally express despair and/or distress, e.g. during restraint (either by jacket, tube, or headpost) and/or cold stress (Grimsley et al., 2016; Yamauchi et al., 2022). However, the established, best-known example for calls expressing a negative affective state in mice are squeaks (or broadband vocalizations, or low frequency harmonics). They are characterized by a fundamental frequency of about 3 kHz, an average of three to five harmonics, often with co-occurring NLP such as subharmonics (Finton et al., 2017; Lupanova and Egorova, 2015; Wang et al., 2008). Mice emit squeaks to express pain, despair and/or distress, for instance during tail snipping, when being bitten (Gourbal et al., 2004; Williams et al., 2008), or during courtship with concomitant defensive behavior (Finton et al., 2017; Lupanova and Egorova, 2015; Sugimoto et al., 2011; Wang et al., 2008). A link between anxiety and squeaks is suggested, as squeaks evoked by a tail-suspension test in mice bred for high anxiety-related behaviors, but not CD-1, BALB/c, DBA, or B6N, are sensitive to diazepam treatment (Ruat et al., 2022).

In contrast, a link between anxiety and LFV emission is not clear. While pharmacological manipulation of anxiogenic neural circuits by the anxiolytic benzodiazepine midazolam or the putative anxiolytic δ opioid receptor agonist KNT-127 causes an overall decrease of calls emitted by animals experiencing restraint and/or cold stress, midazolam has no effect on and KNT-127 even elevates the proportion of LFVs emitted (Yamauchi et al., 2022). However, playing back LFVs to mice that had prior restraint experience leads to an increase in intra-amygdalar acetylcholine concentration, as well as flinching (Ghasemahmad et al., 2024). Cholinergic signaling in the basolateral amygdala has been shown to be important for the durability of fear memory in rats (Crimmins et al., 2023).

In our study, separating same-sex mouse dyads with a divider was suffcient to evoke the predominant emission of LFVs together with Noisy calls. Treatment with the anxiolytic 5-HT_1A_ receptor agonist buspirone reduced the call rate compared to saline treatment, however, the call rate after buspirone treatment was not different from that of completely untreated mice, suggesting buspirone treatment would normalize the effect of the injection rather than alleviating anxiety caused by separation. Furthermore, we did not observe any effect of buspirone treatment on the proportion of LFVs or any other call class.

Squeaks are clearly used by mice to express a negative affective state provoked by another organism. LFVs have, so far, only been reported in mice during considerable restriction of their freedom of movement, suggesting they are also used to express a negative state. However, we have used a gentle separation-method allowing animals at least limited freedom of movement, that is used routinely e.g. for the exposure of naïve male mice to female mice (Klaus et al., 2025; Matsumoto and Okanoya, 2018; Musolf et al., 2010; Zala et al., 2017). Yet, we have observed LFVs to be one of the two dominant (about 40% of all calls) call classes used in this context. Even more so, once direct sensory (and/or tactile) interaction was facilitated with larger holes, LFVS became the dominant call class in the call repertoire with about 75% of all calls. Hence, LFVs might not be limited to the expression of an aversive state, but may encompass a broader communicational role when mice are restricted in their direct interaction, but yet free to move around the test arena.

### Communicating at distance using Noisy calls and LFVs

Until recently, it was agreed upon that mice vocalize mostly at a frequency range from 40 kHz and above with the exception of squeaks (Lupanova and Egorova, 2015; Portfors, 2007; Venkatraman et al., 2024). However, most vocalization studies in adult mice allow them to interact directly with one another. Our data clearly show that preventing this direct interaction leads mice to produce predominantly calls with a low dominant frequency. Considering the low-pass filter properties of the atmosphere (Lawrence and Simmons, 1982), it seems reasonable to use calls with more energy in the low frequency components (Noisy calls and LFVs) as these calls propagate a farther distance than those with more energy in the high frequency components, and also cost less energy to produce. Moreover, mice show the lowest hearing thresholds for tone frequencies ranging from about 10 to 20 kHz (Ohlemiller et al., 2010; C. R. Park et al., 2024). Recently, it was highlighted that both Noisy calls and LFVs (referred to as MFVs in Tehrani et al., 2014) elicit widespread neuronal activity throughout the IC and along its tonotopic axis compared to a more limited neuronal activity in response to USVs (Tehrani et al., 2024). The presentation of noise (non-stationary noise, white noise or natural stationary noise) increases neuronal activity in most sites of the auditory midbrain of both guinea pig and rat (Hosseini et al., 2021; Mishra et al., 2021). However, organization of the auditory midbrain into zones with different response properties as well as the higher energy of non-stationary noise in lower frequencies compared to higher frequencies has to be considered (Chen et al., 2012; Hosseini et al., 2021).

Taken together, the auditory sensitivity of mice in the 10 – 20 kHz frequency range, the wide spread activity along the tonotopic axis in the auditory midbrain, and the putatively lower energetic production cost suggest a key role for Noisy calls and LFVs in the vocal communication of mice when interaction along other sensory modalities is not feasible.

### Few and far between – MFVs and USVs in separation-induced acoustic communication

In all our separation experiments we encountered a small proportion of calls with peak frequencies between 32 and 50 kHz (MFVs). They formed a distinct peak in the peak frequency distribution and their spectrograms appeared tonal and differed from the low frequency component of step USVs. Hence, we regard MFVs as a separate call class. Reports on mouse calls in the MFV peak frequency range are rare, however, so far, they are restricted to mixed-sex interactions. Male mice have been shown to emit calls in the MFV range during ejaculatory mounts (White et al., 1998), while female mice emit 40 kHz calls to initiate pup retrieval in the pup’s father (Liu et al., 2013). Based on the exemplary spectrograms provided, a proportion of copulatory calls would likely be identified to be the low frequency component of step USVs (Arriaga and Jarvis, 2013; Klaus et al., 2025; Scattoni et al., 2008; Wang et al., 2008). The 40 kHz pup retrieval calls likely express an agitated state, as an elevated corticosterone plasma level has been reported in dams that were removed from their litter (Moles et al., 2008). To our knowledge, this is the first time MFVs have been reported to occur during same-sex vocal communication, scarcely, yet consistently. Unfortunately, we were unable to elucidate their role for mouse social behavior any further.

USVs are the best researched class of mouse call, being emitted both during same-sex (de Chaumont et al., 2021; Moles et al., 2007; Panksepp et al., 2007) and mixed-sex interactions (Hanson and Hurley, 2012; Panksepp et al., 2007; Schleidt, 1951; White et al., 1998). Surprisingly, in our study separated mice emitted only few, short USVs of lower peak frequency compared to those emitted during direct same-sex interaction (Caruso et al., 2022; de Chaumont et al., 2021; Ferhat et al., 2016).

## Conclusion

Preventing direct physical interaction of same-sex mouse dyads through a divider revealed a call repertoire consisting mainly of Noisy calls and LFVs. In light of the auditory sensitivity, widespread IC response, and dominant frequencies with greater propagation range, Noisy calls and LFVs appear ideal for communication when e.g., whisking is prevented. In this situation Noisy calls could serve the purpose to attract the attention of other mice, which declines with the concomitant rise of direct interaction possibility. During unrestricted direct interaction (e.g., in a resident-intruder scenario), however, Noisy calls might serve a different role. Until now LFVs have only been described in aversive contexts, however, our study, using a gentle separation design, extends the putative communicational role of LFVs beyond the expression of aversive states.

Taken together, the communication of mice at frequencies below 30 kHz seems to play a much larger role than previously thought, and may even, in certain situations, be the predominant form of mouse vocalisation. As such, research into the vocal communication of mice should extent its scope beyond USVs to obtain a complete picture.

## Materials and Methods

### Animals

Experimental procedures were approved by the State Office for Health and Social Affairs Berlin under the animal license numbers “G0045/22” and “G0276/18”. The FVB.129P2-*Pde6b^+^ Tyr^c-ch^*/AntJ mouse strain (strain no. 004828; Jackson Laboratories, USA) was chosen since these mice do not show retinal degeneration and do not exhibit early onset high frequency hearing loss, making it particularly suitable for behavioral vocalization experiments (Errijgers et al., 2007; Garcia-Pino et al., 2017; Ho et al., 2014). Founding mice were sourced from Jackson Laboratories and bred in-house at the animal facility of Freie Universität Berlin. C57BL/6J (B6J) mice arrived from the Max Rubner laboratory (German Institute of Human Nutrition, Potsdam, Germany). CBA/J (CBA)mice were obtained from the Department of Otolaryngology - Head & Neck Surgery, University of Tübingen Medical Center. Both B6J and CBA mice were allowed to acclimatize to the housing facility for at least 14 days before entering experiments. Mice were housed in standardized home cages (Type II L), with group sizes of two to four individuals per cage. The animals were housed on a 12:12 light-dark cycle (lights on at 3 a.m.) and the facility temperature kept at 22±1°C with relative humidity of 45-65%. Food and water were provided *ad libitum*. The experiments were conducted on mice aged between P52 and P124 (median: P82).

### Experimental Setup

Behavioral experiments were conducted in a custom-built arena with a square area of 930 cm² and a wall height of 30 cm, constructed from Makrolon® and divided into two equally sized compartments by a transparent Plexiglas® wall. No bedding was used during the experiment. The divider featured a perforated section (8.5 cm x 27 cm (HxB)) at the bottom. For the majority of experiments this section featured holes of either 0.5 cm diameter (small holes) or 1 cm diameter (large holes), or was not perforated (no holes). Hence, the two mice could engage with each other using auditory, visual, and olfactory cues, while the exchange of somatosensory cues was severely limited or not possible at all (Hoier et al., 2016; Pessoa et al., 2022). The arena was placed within a sound-attenuated chamber (METRIS-Systems, Hoofddorp, Netherlands) during the experiments. Video recordings were captured at 30 fps using EthoVision XT software (Noldus, Wageningen, Netherlands) and a Brio Ultra HD Pro Webcam (Logitech, Apples, Switzerland) mounted 35 cm above the arena. Audio signals were recorded using a microphone (CM16 Avisoft-Bioacoustics) positioned 30 cm above the arena. The microphone was connected to an CMPA40-5V amplifier (Avisoft Bioacoustics) and the signal was digitized with a sample rate of 384 kHz using an ADI-2-Pro analog-to-digital converter (RME, Haimhausen, Germany) and the Avisoft-RECORDER software (Avisoft Bioacoustics, Glienicke, Germany).

### Experimental procedures

In total 48 pairs of FVB mice (24 male and 24 female pairs), 5 pairs of B6 mice (all male), 5 pairs of CBA mice (2 male and 3 female pairs), and 5 mixed-sex FVB mouse pairs were employed in the physical separation experiment.

All experiments were conducted at dusk (‘lights on’ in the animal facility) to align with the crepuscular activity of mice. Mice underwent a two-day habituation protocol prior to the experiments. On the first day, the animals were transferred to the experimental room for one hour while remaining in their home cages, with food and water provided *ad libitum*. On the second day, the animals were again habituated to the experimental room for one hour before being individually placed into one of the arena compartments for a 20-minute habituation period. After this, they were returned to their home cages.

Before the experiment, the animals were habituated again to the experimental room for one hour in their home cages. Following this, two individuals were transferred to the test arena and placed each in one of the compartments. The arena was then placed inside the sound-attenuated chamber, and video and audio recordings were collected continuously for 15 minutes. After this recording session, the Plexiglas® divider (small holes) separating the two compartments was removed, allowing the mice to physically interact. Video and audio recordings continued for an additional 5 minutes. In the case of dividers without holes, no unification phase was initiated, as the arena contained a lid in order to minimize physical, olfactory and vocal communication between the respective mice.

For the investigation of buspirone’s effect on the separation-induced vocal behavior, the experiments were conducted as described above with the exception that animals received either physiological saline or buspirone hydrochloride (4 mg/kg at 0.1 ml/10g body weight; abcr GmbH, Karlsruhe, Germany, art.no. AB348846) 20 min before the start of the experiment. After the experiment mice returned to their home cage and were transferred back to the animal facility. The experiment was repeated after 3 days, with the same mice now receiving the respectively other substance, be it saline or buspirone. This way the effect of the experimental sequence could be accounted for. A total of 5 male and 5 female same-sex FVB mouse pairs were used for in these experiments.

### Vocalization Analysis

Audio recordings were analyzed -first by creating spectrograms from the original wav files using DeepSqueak 3.1.0, a MATLAB-based platform for detecting and visually illustrating bioacoustic signals (Coffey et al., 2019). Vocalizations were initially detected using the Mouse Detector YOLO2 convolutional neural network, which effectively identifies ultrasonic vocalizations (USVs) but is less sensitive to detect lower-frequency calls. All automatically detected vocalizations were manually reviewed and corrected. Detection boxes were redrawn to precisely match the vocalization contours, ensuring they covered the entire spectral morphology. Vocalizations missed by automatic detection were manually added by drawing detection boxes around the call contours in the spectrogram. For noisy vocalizations without clear spectral morphology, and vocalizations of low frequency (<32 kHz) with multiple harmonics, detection boxes were drawn around the entire signal. To distinguish vocalizations from movement-induced noise, we used multiple verification criteria: presence of tonal call contours in the spectrogram, regular production patterns in bouts similar to previously observed vocal sequences, and audio playback verification using DeepSqueak’s down sampling feature (factor of 20). This way low-frequency vocalizations were additionally verified by having similar acoustic features such as downward and upward sweeps or frequency steps similar to USVs.

For classification, the spectral and temporal parameters (Peak Frequency, Call Length) of the detected vocalizations were exported from DeepSqueak. Bandwidth was extracted by modifying the DeepSqueak CalculateStats.m function to extract the frequency bounds of the user-drawn box surrounding the call contours. Vocalizations were classified based on their peak frequency into Low-Frequency Vocalizations (LFVs; <32 kHz), Middle-Frequency Vocalizations (MFVs; 32–50 kHz), and Ultrasound Vocalizations (USVs; >50 kHz). Noisy vocalizations, were manually classified based on their spectral appearance and large bandwidth.

Smooth example spectrograms were created by converting the recordings through fast Fourier transformation (nfft: 4096 samples, Hamming window: 512 samples, overlap: 256 samples) in MatLab.

### Call spectrogram feature extraction using convolutional neural network (CNN) and clustering

All computational steps were made using Python (v. 3.10.10). Single calls were extracted from recorded audio WAV files using the timestamps obtained by call detection as described above. The calls’ spectrograms (nfft: 256 samples, Hanning window: 256 samples, overlap: 128, linear detrending; matplotlib 3.10.1) were plotted, and the resulting images were rescaled to 224 by 224 pixel size. Default VGG16 image preprocessing was used before feeding the images into the CNN.

We employed a keras (v. 3.9.0) implementation of VGG16 neural network model (Simonyan and Zisserman, 2014) that achieved 92.7% accuracy in the ImageNet Large Scale Visual Recognition Challenge 2014 (ILSVRC2014) outperforming other state of the art CNN models for computer vision such as Clarifai. VGG16’s architecture consists of 13 convolutional layers, 5 max-pooling layers, 3 fully connected layers (fc), and 1 soft-max layer, allowing for extraction of complicated image features. To extract image features from our call spectrograms, we used VGG16 with the weights obtained by previous training on the ImageNet dataset (https://www.image-net.org). A similar approach of clustering images using VGG16 - extracted image features has been used successfully in the past (Cohn and Holm, 2021; Lachmann et al., 2022). To extract features, the network was truncated at the second fully connected layer (fc2), resulting in a 4096-dimensional feature vector for each image. Images were clustered using UMAP (umap-learn library, v 0.5.7) based on the obtained features. To optimize the UMAP projection, a grid search was conducted to identify optimal values for the ‘n_neighbors’ and ‘metric’ parameters. These parameters significantly influence the resulting embedding by controlling the local and global structure of the data. The number of neighbours (5) and Manhattan distance were chosen based on their ability to effectively cluster the data points while preserving the underlying manifold structure, assessed through parameters such as peak frequency, bandwidth and call duration of calls and the visual similarity of spectrograms projected on the embedding. Additionally, the effect of individual animals emitting the call was checked in order to make sure it does not affect the clustering by forming separate, individual-specific clusters.

### Behavioral analysis

Videos were analyzed using DeepLabCut V2.2.1 (Mathis et al., 2018). Nine body parts were defined for each mouse: snout, left ear, right ear, left shoulder, right shoulder, left hip, right hip, tail base, tail tip. The network (dlcrnet_ms5 with multi-animal-imgaug and ellipse tracking method) was retrained on 950 frames from 38 videos each featuring the same setup as for the experimental videos (left mouse = mouse in left compartment, right mouse = mouse in right compartment). Following automatic body part detection, outlier frames were extracted using DeepLabCut’s ‘jump’ algorithm and were corrected manually by adjusting body point markers in the automatically selected frames. The dataset was updated accordingly. The obtained coordinates were filtered and interpolated using SARIMA algorithm (AR: 1, MA: 1, cut-off p: 0.01). The filtered x-y-coordinates were exported from DeepLabCut in the form of a CSV file.

We analyzed the proximity of the snouts of the two mice at the time of call emission (snout-to-snout distance). Snout-to-snout distance was calculated by applying Pythagorean theorem: the length of the direction vector between the snout of the left and of the right mouse were calculated using the x-y-coordinates obtained from DeepLabCut. Then the frame number was translated into time information. The snout-to-snout distance time stamps were then compared to the time stamps marking the beginning of detected vocalizations. Frames with the smallest time difference to call onset were selected for analysis. To determine the mouse body length, the length of the distance vector between snout coordinates of each mouse and the coordinates of their own tail base was calculated.

Furthermore, to visualize whether certain snout-to-snout distances coincided with call production more often than others, we calculated the call probability by dividing the amount of calls at respective snout-to-snout distances by the overall occurrence of this snout-to-snout distance during the experiment. For line plots, data was smoothed using a moving average with a span of 0.025.

Additionally, snout-to-snout angles (during separation phase) as well as snout-to-tail base angles (during unification phase) were calculated as follows, with the head center being defined as the half distance between left and right ear (between-ear-point, B) for each mouse:

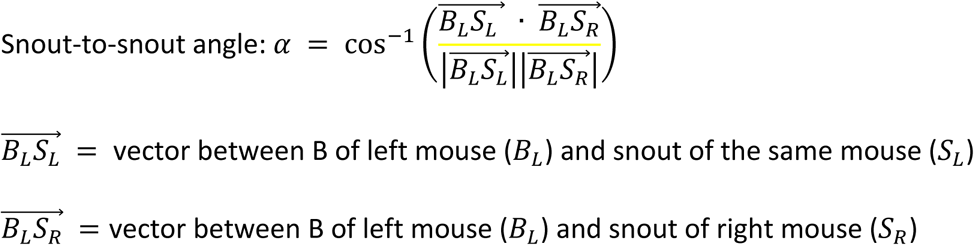

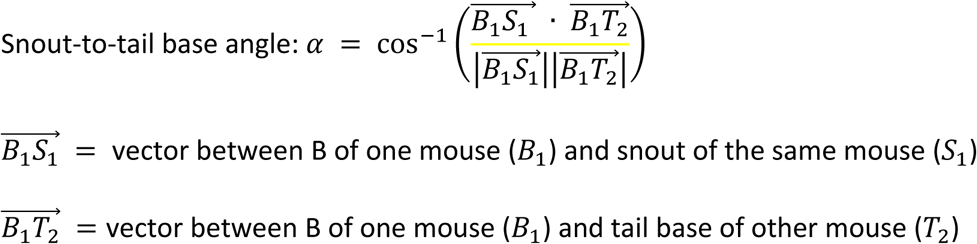

Snout-to-tail base angles were calculated for both the left mouse’ snout to the right mouse’ tail base and the right mouse’ snout to left mouse’ tail base. The smaller of the two angles and its corresponding distance were used for analysis.

### Data Analysis

All data are presented as the mean ± standard deviation of the mean. Data fulfilling the requirements for parametric statistics was analyzed statistically using two-way repeated measures ANOVA (2-way RM ANOVA) for comparisons among three or more groups, with call type as within-subject factor and event as between-subject factor. In the case that 2-way RM ANOVA could not be calculated due to missing values, a restricted maximum likelihood (REML) mixed-effects analysis was employed. If ANOVA results were significant, Sidak’s multiple comparison tests were performed. For comparisons between strains and hole sizes a one-way ANOVA was employed. When parametric statistics requirements were not met Kruskal-Wallis test was employed to examine both strain and hole differences. Sex differences and differences between mice receiving treatment (buspirone or vehicle) and mice not receiving any treatment were analyzed using unpaired t-tests, when requirements for parametric statistics were met. In case the data did not meet these requirements, Mann-Whitney-U test was used for analysis. For comparisons between buspirone and vehicle treatment a paired t-test was used if requirements for parametric statistics were met. If these requirements were not met, Wilcoxon signed rank test was used. Correlations between snout-to-snout distance and angle were analyzed using Spearman correlation analysis. Statistical computations were performed using GraphPad Prism (V. 8, GraphPad Software Inc, U.S.A).

## Supporting information

Archive containing .xlsx files for each figure.

MatLab script used for the generation of line plots in fig. 5

## CRediT authorship contribution statement

**Daniel Breslav:** Conceptualization, Methodology, Data Curation, Investigation, Formal Analysis, Visualization, Writing – Original Draft, Writing – Review & Editing

**Michal Wojcik:** Formal Analysis, Visualization, Writing – Original Draft, Writing – Review & Editing, Funding Acquisition

**Ursula Koch:** Conceptualization, Methodology, Writing – Review & Editing, Supervision, Project Administration, Resources

**Thorsten Becker:** Conceptualization, Methodology, Investigation, Formal Analysis, Data Curation, Visualization, Writing – Original Draft, Writing – Review & Editing, Supervision, Project Administration, Funding Acquisition

## Acknowledgements

We thank Virginia M. Baatz, Johanna M. Kube, Julia Freitag, and Luna Reimer for assistance with experimentation. This work was supported by the German Center for the Protection of Laboratory Animals (Bf3R) under award number 60-0102-01.P615. M. Wojcik was awarded the Elsa Neumann Scholarship.

## Data availability statement

All single calls and their spectrograms are deposited at https://zenodo.org/records/17581239. All data used for mouse-to-mouse distances and body angles are deposited at https://zenodo.org/records/17491957. All data used for VGG16-feature extraction from spectrograms and UMAP are deposited at https://zenodo.org/records/17493153. Data is publicly available.

The code for VGG16-feature extraction is available at https://github.com/tmb404/MouseCallClustering. The code for analysis of mouse-to-mouse distances and body angles is available at https://github.com/tmb404/MouseBehav.

All other data generated or analyzed during this study are included in the manuscript and supporting files; source data and code files have been provided for all figures.

---

**Fig. 4 – Suppl. 1.**
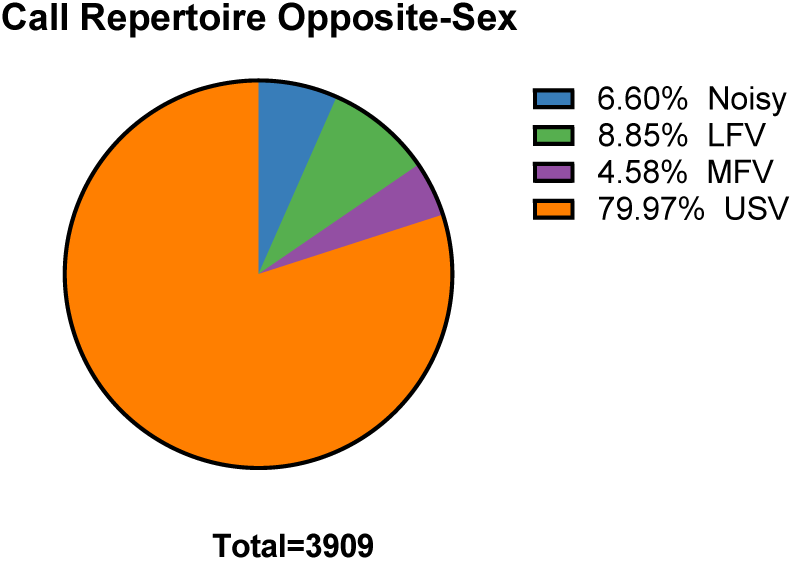

**Fig. 8 – Suppl. 1.**
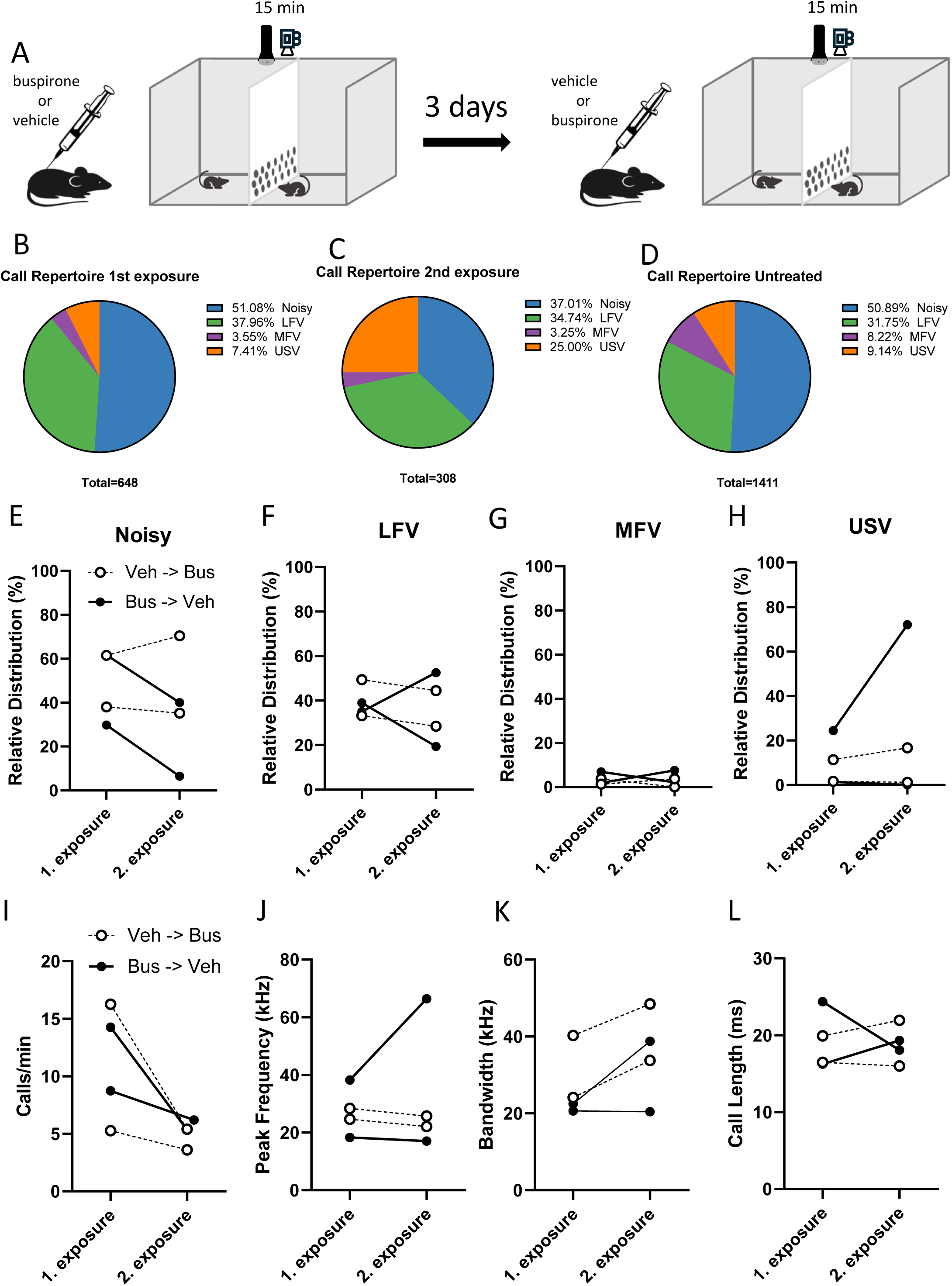

